# The Macerprep: a minimalist kit- and enzyme-free high-yield miniprep utilising alkaline lysis and alkaline hydrolysis principles

**DOI:** 10.1101/2020.08.13.249607

**Authors:** Ian Hu

## Abstract

The commercialisation of miniprep kits supplanted the original alkaline lysis method for plasmid DNA preparation, and had remained relatively unchanged for almost two decades. The Miraprep substantially improved the yields of miniprep kits. However, the method still relies on commercial kits, which can be a burden financially to certain projects. Additionally, Pronobis et al. also identified loss of RNAse activities in miniprep kits over time. The present novel plasmid DNA isolation protocol addresses the two issues mentioned above utilising alkaline lysis and alkaline hydrolysis principles. With a largely identical workflow and operation time, the Macerprep will significantly reduce costs of establishing new laboratories as well as maintenance of running molecular biology laboratories.

## INTRODUCTION

Plasmid purification from *Escherichia coli* is one of the most used tools in a molecular biologist’s arsenal. The alkaline lysis method (Birnboim & Doly, 1979) has always been the preferred method since its conception –although there are few alternatives (Ouyang *et al*., 2008)– and even as commercial kits superseded the traditional method, the basic principles of alkaline lysis and neutralisation remains (Colpan *et al*., 2002). While the commercial miniprep kits brought the handling and harvesting time down from half a day to less than 30 minutes, the yield, however, also reduced substantially, spurring the emergence of larger formats of silica column midi-, maxi-, and mega-prep kits, which seem to be priced according to their yields. The introduction of the Miraprep has changed this; by the simple addition of ethanol before the silica-binding step, the yield of a miniprep column is improved 10-fold (Pronobis, Deuitch & Peifer, 2016).

The principle which causes the DNA to bind with silica seems to be related with DNA condensation. Under chaotropic conditions, the hydrogen bond between the DNA molecules and water is disrupted, forcing DNA to bind to the silica, or further, collapse onto itself (Boom *et al*., 1990; Bloomfield, 1997; Vandeventer *et al*., 2012; Green & Sambrook, 2017; Turner *et al*., 2018; Hu, 2019). In the case of column purification, the DNA is then washed with an ethanol-based buffer which maintains the low DNA solubility and removes the chaotropic salts, and finally eluted with small amount of aqueous buffer to obtain a highly concentrated DNA sample (Qiagen, 2020a, 2020b). Most commonly, the chaotropic salt used in a commercial kit is guanidium chloride. In Qiagen’s original patent, a Solution N3, composed of 4.2 M guanidine hydrochloride, 0.9 M potassium acetate, pH 4.8, is described (Colpan *et al*., 2002), and is suspected to be used in most, if not all, miniprep kits.

To remove the substantial amount of bacterial RNA in the prep, RNAse is routinely employed. However, the RNAse has a tendency to lose activity over time (Pronobis et al.). The Miraprep protocol recommends adding fresh RNAse to the resuspension solution before each purification. This method is certainly doable and practicable, yet increases logistic burden of the routine lab maintenance. The present study ventures utilising the principle of RNA alkaline hydrolysis as an effective method to improve DNA purity and yield. RNA is notoriously unstable and prone to degradation due to the extra hydroxyl group on the 2’ position in the ribose when compared with DNA. The hydroxyl group deprotonates and attacks the adjacent phosphodiester bond of the backbone under alkaline conditions, of which the reaction rate is elevated at higher temperatures (Coward, 2012). The phenomenon is most often undesired, whereas its only usage in biology until now seems to be short RNA probe production for *in situ* hybridisation (Yang *et al*., 1999) and in rare cases, removing dangerous viral RNA from clinical samples (Lemire, Rodriguez & McIntosh, 2016).

The original Miraprep study proposes that one volume of ethanol be mixed with the lysate before passing through the column. Although higher volume of ethanol does seem to improve yield slightly, it also seems to cause more RNA contamination (Pronobis et al.). The alcohol of choice in this study, on the other hand, is isopropanol. For common DNA precipitation with sodium acetate, alcohol usages for ethanol and isopropanol are roughly 2.5-times and 1-time volume of the DNA solution, respectively (Green & Sambrook, 2017). If the two processes, Miraprep/Macerprep and DNA precipitation, share some common biochemical principles, using isopropanol could reduce the total volume of liquid to pass through the silica columns, further reducing operation time. As this study aims to remove RNA with chemical means, a higher percentage of the alcohol with higher chaotropic properties and lower DNA solubility could be beneficial for the final DNA yield.

## RESULTS

### Addition of isopropanol to cleared lysate by traditional method facilitates DNA binding with silica column

Firstly it was tested whether the addition of isopropanol to the cleared lysate from the traditional alkaline lysis method is sufficient for DNA binding with the silica column. 5 mL of *E. coli* strain Dh10 hosting the plasmid pUC19 was incubated overnight with orbital shaking at 37°C. 1 mL of the grown cell suspension was pipetted into a 1.7 mL microfuge tube and pelleted with 1000 rcf at room temperature for 5 minutes. After the *E. coli* pellet was resuspended with 250 *µ*L of Solution 1, cells were broken with 250 *µ*L of Solution 2, and neutralised with 250 *µ*L of Solution 3 and incubated for 5 minutes on ice. The lysate was centrifuged for 20 minutes at 16060 g at 4°C. The cleared lysate before and after the addition of isopropanol was passed through a column, washed twice with 700 *µ*L of Wash buffer, eluted with 50 *µ*L elution buffer, and analysed with agarose electrophoresis. For comparison purposes, another set of lysis reaction neutralised with a homemade Buffer N3 was also included (Fig. 1). The results demonstrate that the cell lysate by the traditional Solution 3 process has virtually no binding with the silica column, and isopropanol does improve DNA binding for this method, yet a significant amount of smaller molecular weight RNA is also retained. Interestingly, for the Solution N3 process, which includes both acetate and guanidium, the addition of isopropanol does not seem to have an effect on the amount of both DNA and RNA in the eluate (Fig. 1).

**Figure 1:**
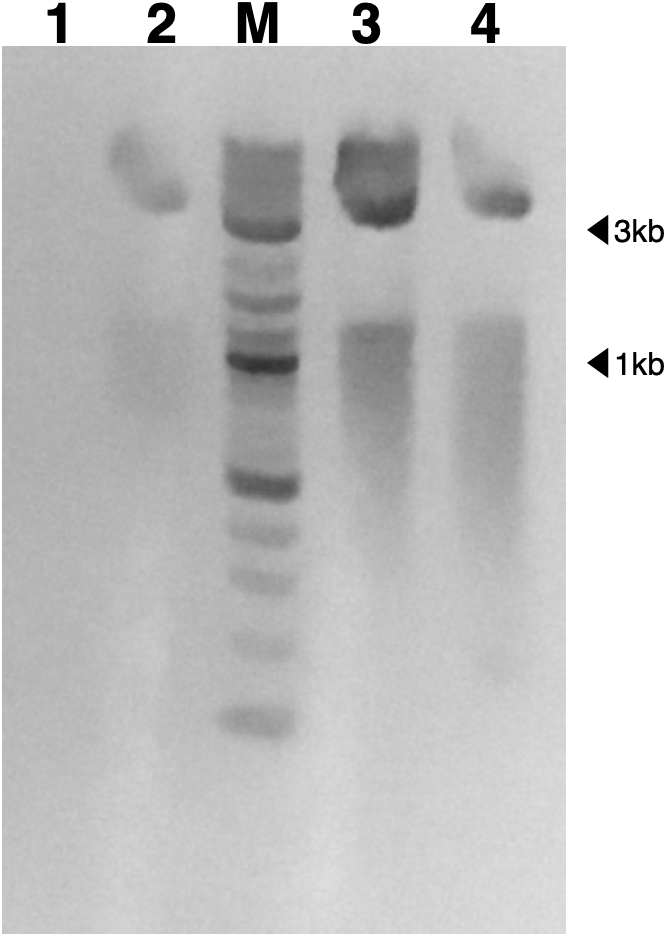
Isopropanol promotes DNA and RNA binding to silica column in the absence of guanidium. Lane 1, 2, column eluate of Solution 3 process before and after addition of isopropanol. Lane 3, 4, column eluate of Solution N3 process before and after addition of isopropanol. M, marker.

### Prolonged alkaline heat treatment greatly reduces RNA and improves plasmid DNA yield

In order to evaluate whether alkaline hydrolysis can be used to remove contaminating RNA without significantly damaging plasmid DNA and reducing yield, the cell lysate after the addition of the alkaline Solution 2 was incubated at high temperature for increasing amount of time. Immediately after the addition of 250 *µ*L of Solution 2 and several inversion, lysed *E. coli* cells in microfuge tubes were then placed on a 95°C heat block for a duration from 0 to 30 minutes. The tubes were then water-cooled to room temperature, the rest of the plasmid prep procedure carried out, and the final eluate were analysed by agarose electrophoresis (Fig. 2). The results clearly demonstrate a reduction of RNA over time of heat treatment, and an increased DNA to RNA ratio. The results show the same trend for both Solution 3 and N3 processes. It was, however, observed that after 10 minutes of heat treatment of both sets of experiments, the integrity of DNA seems to start to be compromised, as a larger molecular weight new species can be observed on the gel. Consequently it was decided that 10 minutes would be the best incubation time, as this is the least amount of time required to diminish the amount of RNA to the level of being unobservable by the staining method utilised while the level of harvested DNA still seems unharmed.

**Figure 2:**
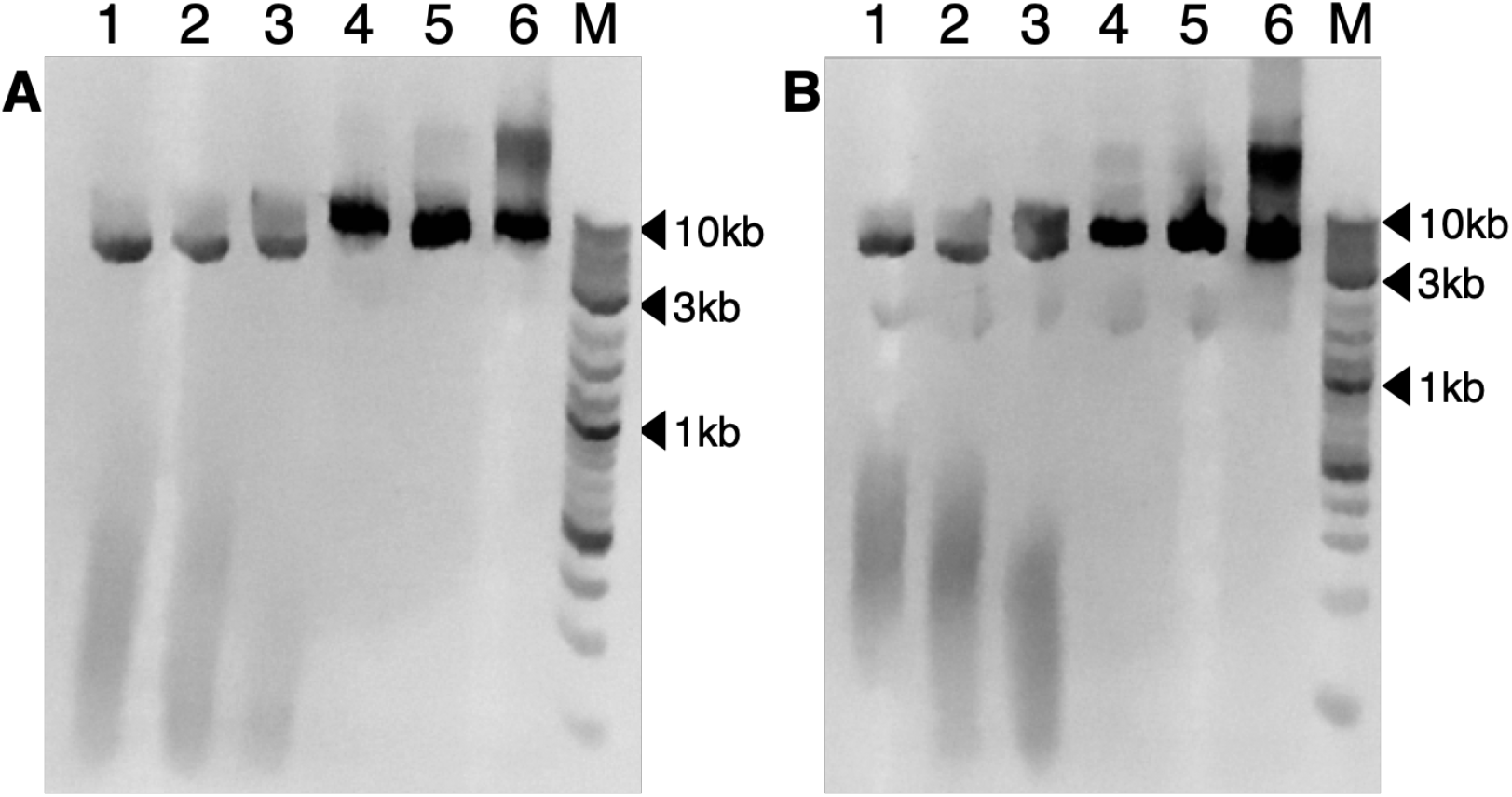
Prolonged heat treament reduces RNA and increases DNA binding to silica column. Bacterial alkaline Iysate were incubated at 95 ^°^ for various amount of time before the addition of either neutralisation buffer. 2A, Solution 3 process. 2B, Solution N3 process. 1, 2, 3, 4, 5, 6, incubation time of 0, 1, 2, 5, 10, and 30 minutes at 95 ^°^. M, marker.

### The Macerprep performs similarly when compared with the Miraprep

To compare the results from a commercial kit using the Miraprep protocol with the newly developed Macerprep procedure, column eluates from both procedures were run alongside each other on an agarose gel (Fig. 3). The result suggests that there are no significant differences between the DNA preps obtained from both methods, either in terms of yield or level of RNA contamination.

**Figure 3:**
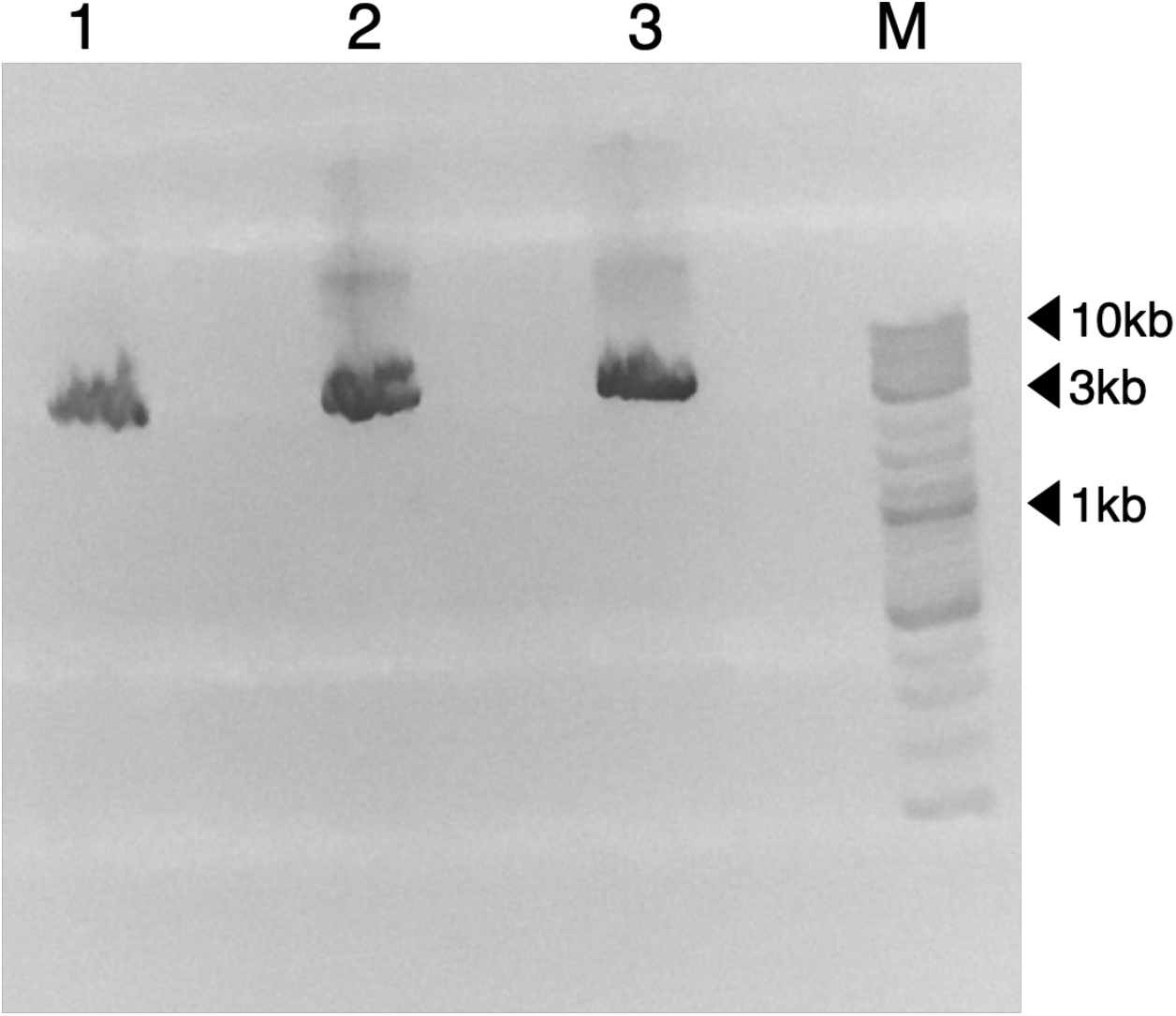
No obvious differences are observed between the Miraprep and the Macerprep. 1, the Miraprep. 2, the Maceerprep with Solution 3 process. 3, the Macerprep with Solution N3 process.

To further characterise the usability of the Macerprep, the plasmid DNA was then subjected to a restriction enzyme digestion, PCR reaction, and direct Sanger sequencing. Macerprepped DNA along with Miraprepped DNA were digested by the restriction enzyme DraI (Fig. 4). The result demonstrates that all three DNA prep were sufficiently digested.

**Figure 4:**
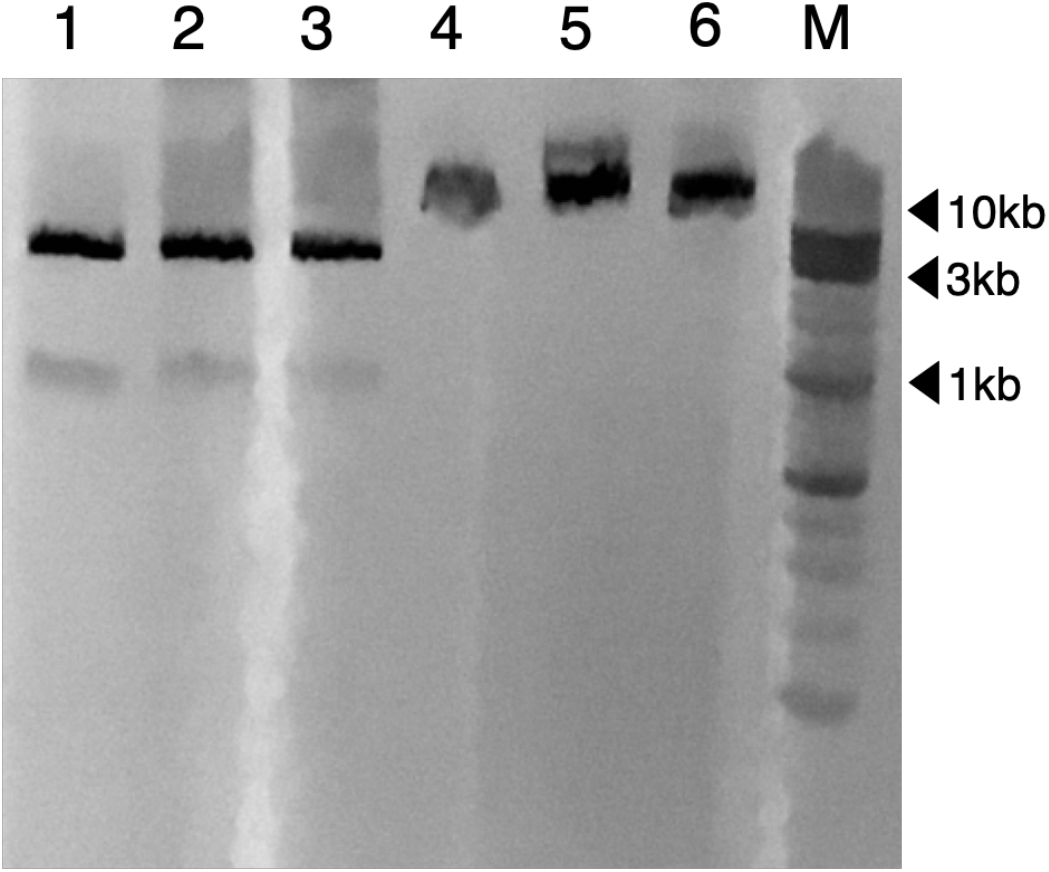
Macerprepped DNA is good template for restriction enzyme. 1-3, post Dral enzyme digestion. 4-6, before enzyme digestion. 1, 4, Miraprepped DNA. 2, 5, Macerprepped DNA with Solution 3 process. 3, 6, Macerprepped DNA with Solution N3 process. M, marker.

For the restriction enzyme test, the preps were then diluted 100 times in volume to adjust to a concentration suitable for PCR, and were used as templates for two polymerases, the NEB Q5 HiFi polymerase and NEB Onetaq 2X master mix, with primer sets designed against the beta lactamase gene. All combinations show a single specific PCR product at the expected size (Fig. 5).

**Figure 5:**
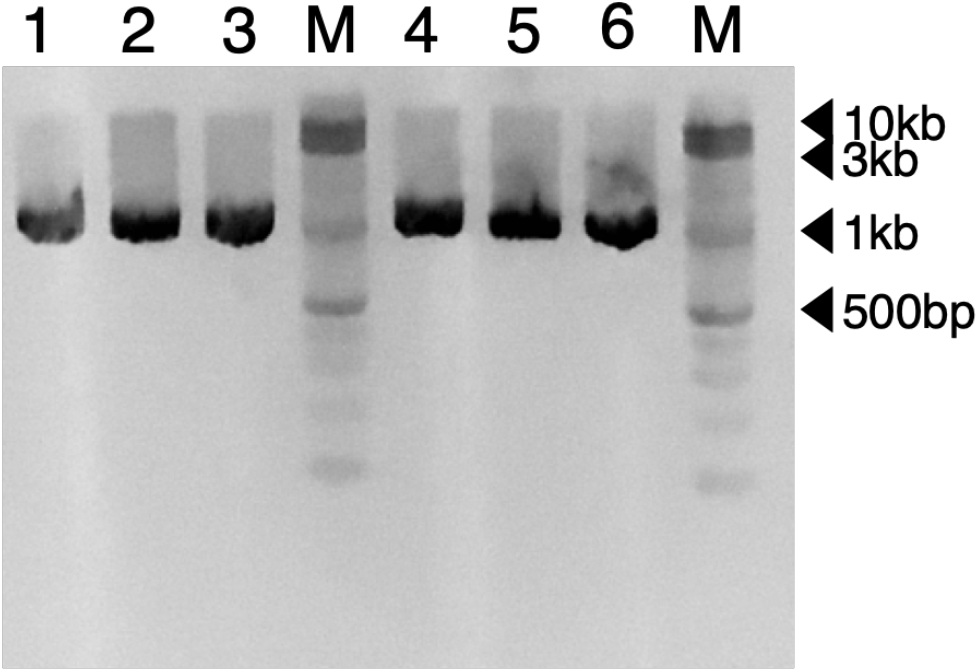
Macerprepped DNA is good template for PCR. 1-3, PCR product by NEB Q5® HiFi polymerase. 4-6, PCR product by NEB Onetaq polymerase. I, 4, Miraprepped DNA. 2, 5, Macerprepped DNA with Solution 3 process. 3, 6, Macerprepped DNA with Solution N3 process. M, marker.

FInally, direct Sanger sequencing of all three preps demonstrate satisfactory results. To mimic common laboratory malpractice and test the stability of the prepped DNA, all three preps were incubated at room temperature for 96 hours before subjecting to Sanger sequencing. All preps provided good sequencing results with close to or above 1000 bp of easily legible chromatograms. An excerpt of the alignment of three chromatograms is shown (Fig. 6). No conspicuous differences can be observed between the three in terms of chromatogram quality and signal strength.

**Figure 6:**
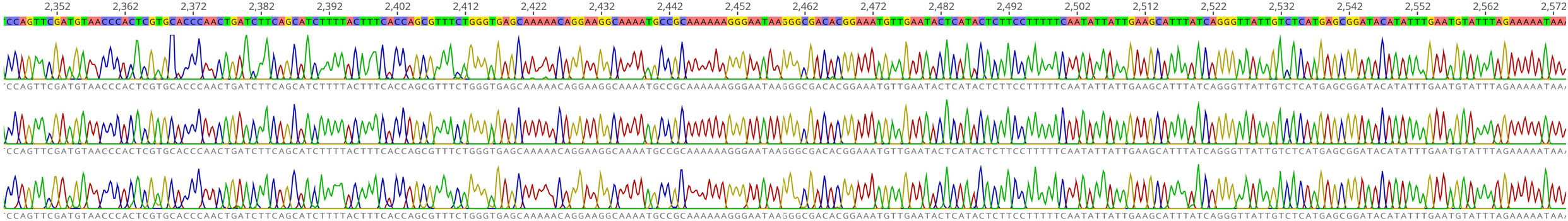
Macerprepped DNA can be directly Sanger sequenced. Top row, raw chromatogram of Miraprepped DNA using Thermofisher GeneJet miniprep kit. Middle and bottom row, Macerprepped DNA with Solution 3 and N3 processes.

## DISCUSSION

The final Macerprep protocol utilises a minimalist approach; the cell resuspension solution contains only sodium chloride and EDTA with no additional sugar or lysozyme, and the neutralisation solution with only sodium acetate, pH adjusted by HCl. The sodium chloride/EDTA combination is routinely used for *E. coli* protein expression workflows (Turner *et al*., 2018; Hu, 2019) and should be sufficient in keeping the bacteria alive shortly; although the difference in osmotic pressures between the growth media and the resuspension buffer may cause harm to the cells long term-wise, the buffer only needs to maintain the cells for until the lysis buffer is added, and the exclusion of sugars and RNAse further prevents the solution from spoiling and microbial growth and negates the necessity to store the solution refrigerated. The neutralisation solution described in Qiagen’s patent (Colpan *et al*., 2002) is composed of potassium acetate and guanidium chloride. Although the potassium salt has the benefit of forming the insoluble salt with the dodecyl sulfate ion, chilling achieves largely the same effect of greatly reducing the solubility of sodium dodecyl sulfate, and the two wash steps incorporated in this method seem to be enough for the final plasmid DNA for enzyme digestion, PCR, and direct Sanger sequencing. Additionally, as demonstrated by the results provided in this study, the inclusion of an additional chaotrophic salt, guanidium chloride, is not necessary, as isopropanol is enough to promote DNA-silica binding, or even DNA precipitation at the atomic level, demonstrated in Pronobis et al. with filter papers. This neutralisation solution also shares its sole component with the commonly used alcohol DNA precipitation protocol, further removing the need to acquire and store an extra chemical whose usage is scarce outside of DNA purification.

It is observed that the cleared crude lysate obtained by the traditional method alone does not provide the affinity of DNA and silica columns. The addition of isopropanol caused both plasmid DNA and RNA to bind to the columns, presumably competing for the binding capacity. On the other hand, the inclusion of guanidium salt in the neutralisation solution provides enough affinity for the nucleic acids and the column and allows for the binding of both RNA and DNA and their competition for the column to occur, yet the addition of isopropanol did not increase the yield of either DNA nor RNA. This was somewhat surprising: Pronobis et al. proposed that in the Miraprep, the columns act partly as a particle filter for the alcohol-precipitated DNA. If such precipitation process had occurred in the experiments conducted for Fig. 1, one would expect the DNA yield to still increase simply by the very same filtering effect in spite of the presence of RNA. This however was not the case. On the other hand, alcohol precipitation of plasmid DNA has been a trial-and-tested method for decades, and is known to precipitate, given enough time, both plasmid DNA and RNA [traditional method]. There are two potential possibilities among others for such contradicting results: the presence of RNA interferes with the time required for the alcohol-precipitation effect, or another novel phenomenon is actually responsible for the fast aggregation of plasmid DNA, which is interfered by the presence of RNA, e.g. crystallisation. These confounding results warrant further studies.

There is concern that the prolonged alkaline heat treatment would cause undesirable effects on the plasmid DNA. Indeed, early pioneering researches in the DNA field suggest that at pH >13, plasmid DNA undergoes an irreversible denaturing process (Pouwels *et al*., 1968; Pouwels, van Rotterdam & Cohen, 1969; Birnboim & Doly, 1979), and that the irreversibly denatured DNA has a higher mobility than its native counterpart. In the present study, no smaller molecular weight species were detected, and if any a slight decrease in mobility seems to be present after prolonged heat treatment, signifying potential plasmid DNA damage. Since electrophoretic mobility is affected by many factors, it is difficult to pinpoint the cause of this phenomenon without further and deeper studies, but the removal of RNA, the ensuing additional binding of plasmid DNA, and the overloading effect of the plasmid band could be one of the culprit. The restriction enzyme tests and Sanger sequencing results support that the plasmid DNA obtained by the Macerprep is of sufficient quality, and no significant differences in quality is observed between Miraprepped and Macerprepped DNA. This study seems to demonstrate that double-stranded plasmid DNA, or DNA in general, is not as vulnerable as we think conventionally. Although no eukaryotic transfection is demonstrated in this study, the author is hopeful that Macerprepped DNA is also suitable for this purpose.

From a financial perspective, an underfunded or proof-of-concept project, whether in academia or industry, often either cannot justify certain spending or simply do not have the budget to spare for high cost products. The costs of commercial miniprep kits from top shelf vendors are not trivial. Third-party silica spin columns, however, can be obtained for a relatively modest price; as an example, for the present study, the columns were acquired for roughly 35 British pence each. As all other components required by Macerprep are all essential chemicals in virtually any molecular biology experiments, the Macerprep truly reduces the cost of plasmid purification by a significant margin. The author wishes this newly developed method will prove useful for and aid new startups or novel studies that has not received grant funding yet, as well as for education purposes.

## MATERIAL AND METHODS

### Macerprep protocol

Bacterial strain and plasmid used for this study are DH10 and pUC19. 3-5 mL of bacterial culture from a single colony on a previously grown LB plate is incubated overnight in a shaking incubator at 37°C. The next morning, 1 mL of the culture was transferred to a 1.7 mL microfuge, pelleted at 4000 rpm for 5 minutes with an Eppendorf 5401 centrifuge. The supernatant was pour off and the pellet was resuspended with 250 *µ*L of Solution 1 (10 mM Tris pH 8, 50 mM EDTA). Afterwards, 250 *µ*L of Solution 2 (1% SDS, 0.2 N NaOH) was added. The tube was reversed several times and then placed on a dry bath set to 95°C for 10 minutes, then cooled with a water bath at room temperature. 250 *µ*L of Solution 3 (3 M sodium acetate, pH adjusted to 5 by HCl) was then added, the tube reversed several times, and then placed on ice for 5 minutes. Alternatively, Buffer N3 (4.2 M guanidium chloride, 0.9 M sodium acetate, pH 4.9) was used. The tube was then centrifuged at 16060 g at 4°C for 20 minutes. 700 *µ*L of the supernatant was carefully removed to a new tube with a micropipette, mixed with 1X volume of isopropanol, and passed through a silica column with a collection tube (Bio Basic inc.) in two batches. The column was then washed twice with the Wash buffer (80% ethanol, 10 mM Tris pH 8), all flow-through decanted after each centrifugation step, and finally centrifuged at 16060 g at room temperature empty for 3 minutes to remove the ethanol. The tube was then placed on the dry block set at 60°C until all the ethanol was dried off. The column was placed inside a new tube, 50 *µ*L of pre-warmed 10mM Tris pH 8 was added to the centre of the column, the column was incubated at 60°C for 1 minute, the DNA was eluted with centrifugation at 16060 g for 1 minute.

### Miniprep kit

When referred to, the Thermofisher GeneJet® miniprep purification kit was used according to the manual with the exception of addition of ethanol according to Pronobis et al.

### Agarose gel electrophoresis and gel imaging

All electrophoresis were performed on 0.9% agarose gel in 1X SAB (10 mM sodium acetate, 10 mM boric acid, pH adjusted to 9), 180 Volts for 20 minutes. Samples were premixed with 3.33X GelRed (Biotium) and 1X NEB purple loading buffer (NEB) before loaded in the wells. After electrophoresis, the gel was imaged on a UV transilluminator (Syngene) and a mobile imaging darkroom with an orange filter, built in-house (Team OpenCell, unpublished), and images taken with an iPhone SE, first edition. Images were then adjusted to black and white and then inverted with the computer software Pixelmator version 3.8.1.

### Polymerase Chain Reaction, restriction enzyme digestion, and Sanger sequencing

PCR was performed with a G-Storm GS482 thermal cycler. The primer set has the following sequence: 5’-CCGCTCATGAGACAATAACC-3’, and 5’-GGTCTGACAGTTACCAATGC-3’. According to the NEB calculator (NEB, online tool), Tm for NEB Q5 and NEB Onetaq was determined to be 52°C and 61°C. Expected PCR product size is 928 bp, and extension was determined to be 50 seconds at 72°C. A total of 30 cycles were performed.

The enzym DraI (NEB) was used for the digestion test. Each reaction was 50 *µ*L in volume consisting of 10 *µ*L of undiluted plasmid prep, 1X Smart buffer, and 10 units of DraI. The reactions were incubated at 37°C overnight.

Sanger sequencing was performed by Genewiz UK with M13 reverse universal primers. Chromatograms were visualised with Geneious Prime 2020.0.5 (https://www.geneious.com).

## ACKNOWLEDGEMENTS

The author sincerely thanks Dr John Waite for his support. The author also wishes to express gratitude towards the team of Better Dairy ltd, as well as the team of the hosting institution Open Cell.

